# DE-Rigor-Agent: An Agentic Workflow for Assumption-Aware Differential Expression Analysis of Bulk RNA-seq Data

**DOI:** 10.64898/2026.11.09.765522

**Authors:** Ziqi Rong, Changhu Wang, Chenxin Flora Jiang, Yiwen Yang, Jingyi Jessica Li

## Abstract

Differential expression (DE) methods for bulk RNA-seq differ in their statistical assumptions, input data requirements, and supported study designs. Large language model (LLM) agents can automate DE analysis, but they may default to popular methods without checking whether these methods are appropriate for the data. We introduce DE-Rigor-Agent, an assumption-aware agent that organizes DE analysis into three stages: input standardization, method filtering, and method execution. Before executing any DE method, the agent uses expert-curated criteria and dataset-specific diagnostics to evaluate six commonly used methods, documents the reasons for retaining or excluding each method, and executes all retained methods. We evaluated DE-Rigor-Agent on 140 simulated datasets spanning 14 challenging scenarios involving small sample sizes, zero inflation, expression outliers, pre-normalized inputs, and confounded batch effects. Benchmarks showed that method-dataset incompatibilities could lead to inflated false discovery rates (FDRs) or reduced statistical power. Two general-purpose LLM agent baselines, including one explicitly prompted to consider statistical assumptions, selected DESeq2 or edgeR for nearly all datasets. In contrast, DE-Rigor-Agent’s filtering decisions closely matched the incompatibilities introduced in the simulations and remained consistent across random seeds, with little additional computational cost. These results demonstrate that DE-Rigor-Agent provides an automated approach to assessing method suitability, making DE analysis more rigorous and transparent.

## 1 Introduction

Differential expression (DE) analysis is a core step in bulk RNA-seq studies for identifying genes whose expression differs between biological conditions [1, 2]. A key statistical challenge is to detect true expression differences while controlling the false discovery rate (FDR). Commonly used DE methods include DESeq2 [3], edgeR [4], limma-voom [5], standard limma without voom [6], the Wilcoxon rank-sum test [7], and NOISeq [8].

DE methods differ in their statistical assumptions and ability to accommodate complex study designs. DESeq2 and edgeR model RNA-seq counts using negative binomial distributions, whereas limma-voom estimates the mean–variance relationship to obtain precision weights for Gaussian linear models. Standard limma applies Gaussian linear models directly to appropriately normalized and transformed expression data. Nonparametric alternatives include the Wilcoxon rank-sum test, which compares expression distributions without assuming a specific parametric model, and NOISeq, which estimates an empirical noise distribution from the data. These methods also differ in their ability to account for covariates. Regression-based methods can adjust for batch effects and other covariates, whereas the standard Wilcoxon rank-sum test does not directly support covariate adjustment.

These differences mean that no single DE method performs consistently well across all datasets [9, 10, 11]. When a method’s assumptions are poorly matched to the data, its FDR control or statistical power may be compromised. For example, in human cohort studies, heterogeneity across individuals can lead to deviations from negative binomial assumptions [12], resulting in inflated FDR for DE-Seq2 and edgeR as sample size (the number of biological samples) increases [13]. In these settings, the Wilcoxon rank-sum test can provide better FDR control but may have limited power at small sample sizes [11, 13]. Benchmark studies further demonstrate that DE performance varies with sample size, outliers, and distributional skew [10, 11, 13]. However, these studies evaluate specific datasets and simulation settings, and their findings may not generalize to a new dataset. Selecting an appropriate DE method therefore requires considering dataset-specific characteristics, including sample size, excess zeros, outliers, input data transformations, and experimental covariates, in relation to each method’s assumptions and requirements.

In practice, assessing whether a DE method is suitable for a given dataset is challenging. It requires translating statistical assumptions and method-specific requirements into concrete data diagnostics, which often demands substantial statistical expertise. Consequently, analysts may rely on familiar methods or default pipelines rather than systematically evaluating method suitability. For example, widely used methods such as DESeq2 and edgeR may be applied without examining whether their distributional assumptions are appropriate for the data. As a result, studies may report lists of DE genes without a clear justification for method selection, even though different methods can yield substantially different results.

The same challenge extends to large language model (LLM) agents, which can plan analyses, write and execute code, and inspect results in response to natural language requests. General coding assistants and specialized biomedical agents have demonstrated the ability to perform complex bioinformatics analyses [14, 15, 16, 17, 18, 19, 20, 21]. However, successful code execution does not guarantee appropriate statistical method selection. As we demonstrate in this study, LLM agents tend to select widely used DE methods without systematically checking whether their assumptions are appropriate for the data. Simply prompting agents to consider these assumptions does not adequately address this problem. These findings motivate incorporating explicit statistical criteria and dataset-specific diagnostics into agentic workflows to support rigorous and transparent DE method selection.

Here, we introduce DE-Rigor-Agent, an assumption-aware LLM agent for bulk RNA-seq differ-ential expression analysis. The agent framework organizes the analysis into three sequential stages: input standardization, method filtering, and method execution. Before executing any DE method, the agent evaluates candidate methods against their statistical assumptions and input requirements using dataset-specific diagnostics. Rather than selecting a single optimal method, it excludes methods that fail these checks and retains those suitable for further analysis. Dedicated subagents then execute the retained methods using expert-curated code templates, producing DE results, an audit log of filtering decisions, and files for reproducibility. By separating method filtering from execution, DE-Rigor-Agent makes method selection transparent and avoids choosing methods based on their observed DE results.

We evaluated DE-Rigor-Agent using 140 simulated datasets spanning 14 scenarios designed to challenge DE methods through small sample sizes, zero inflation, extreme expression outliers, pre-normalized inputs, and confounded batch effects. As a reference, we applied individual DE methods to these datasets and evaluated their FDR control and statistical power. These analyses demonstrated that methods applied to unsuitable datasets could exhibit inflated FDR or reduced power, providing a reference for assessing the agent’s filtering decisions.

We compared DE-Rigor-Agent with two standard Codex agents under different prompting strategies: one was given only the DE analysis task, and the other was additionally prompted to consider method statistical assumptions. All agents were powered by the same underlying LLM. Both Codex agent baselines selected DESeq2 or edgeR for nearly all datasets, even when their assumptions were violated. In contrast, DE-Rigor-Agent systematically evaluated candidate methods and excluded those with detected violations of their assumptions or input requirements. Its filtering decisions closely aligned with the incompatibilities introduced in the simulation scenarios and remained consistent across random seeds, with minimal additional computational cost. These results demonstrate that incorporating expert-curated statistical criteria and dataset-specific diagnostics into LLM agent workflows can improve the rigor and transparency of automated DE analysis.

## 2 Results

### 2.1 Overview of DE-Rigor-Agent

DE-Rigor-Agent organizes bulk RNA-seq DE analysis into three sequential stages: input standardization, method filtering, and method execution (Figure 1). Unlike general-purpose LLM agents that may generate an analysis plan without explicitly evaluating method suitability, DE-Rigor-Agent requires every analysis to pass through method filtering before any DE method is executed. Within each stage, the agent writes and executes code in a preconfigured computational environment and produces a structured report that documents its outputs and decisions. These reports provide a consistent interface between stages and maintain a record of the analysis.

**Figure 1:**
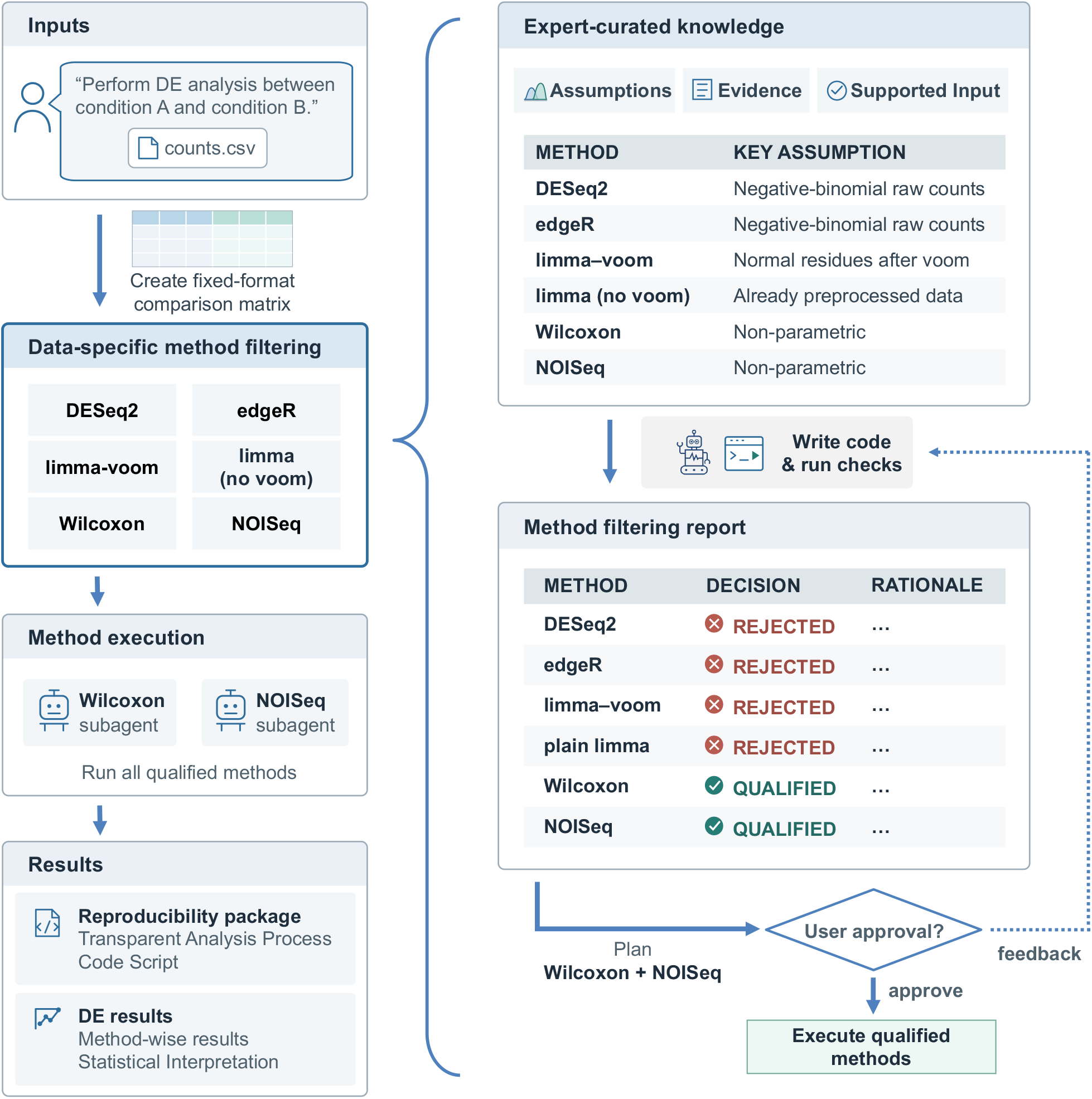
Overview of DE-Rigor-Agent with three sequential stages. In input standardization, the agent converts user-provided files into an expression matrix and sample metadata table. In method filtering, the agent uses expert-curated criteria and dataset-specific diagnostics to evaluate six candidate DE methods, producing a report that documents the decision and supporting evidence for each method. The user can approve the results or provide feedback for reassessment. In method execution, dedicated subagents execute all retained methods with expert-curated code templates. The final output includes DE results, a record of filtering decisions, and files for reproducibility.

#### Input standardization

The first stage converts user-provided files into standardized inputs for downstream analysis. Because input files may differ in format, gene and sample identifiers, and metadata organization, the agent uses the LLM’s general reasoning capabilities to interpret the files and user instructions. It produces an expression matrix with genes as rows and samples as columns, together with a sample metadata table containing the conditions of interest and available covariates. When essential information, such as whether expression values have already been normalized or transformed, cannot be determined from the inputs, the agent requests clarification from the user. Subsequent stages operate on these standardized files.

#### Method filtering

The second stage evaluates six candidate DE methods: DESeq2, edgeR, limma-voom, standard limma without voom, the Wilcoxon rank-sum test, and NOISeq. Rather than selecting a single optimal method, the agent identifies which methods are suitable for the dataset based on expert-curated statistical criteria and dataset-specific diagnostics. The expert knowledge consists of two components (Figure 1). First, method cards specify each method’s distributional assumptions, input data requirements (e.g., raw counts or normalized and transformed expression values), and supported study designs. Second, filtering principles provide additional guidance for specific data characteristics, such as sample size. Guided by these criteria, the agent examines the expression matrix and sample metadata, generates and executes diagnostic code, and produces a filtering report documenting the decision and supporting evidence for each method. The user can approve the filtering results or provide feedback for reassessment, but cannot override the predefined filtering criteria.

#### Method execution

The final stage executes all methods retained after filtering. Each method is assigned to a dedicated subagent, a separate LLM instance equipped with method-specific instructions and code-execution tools. These instructions include an expert-curated, executable code template that the subagent adapts to the dataset by specifying the conditions to compare, incorporating relevant covariates, and performing any required preprocessing. This design allows the agent to accommodate different datasets while maintaining consistent implementations of the retained methods. The main agent then consolidates the results into a final report that summarizes and compares the identified DE genes across methods, documents the filtering decisions, and provides the analysis code and execution records for reproducibility.

### 2.2 Simulation Design and DE Method Performance

To demonstrate the consequences of applying DE methods without assessing their suitability for the data, we evaluated the FDR and statistical power of six commonly used DE methods across simulated datasets with different characteristics. We designed scenarios involving small sample sizes, deviations from distributional assumptions, incompatible input transformations, and confounded batch effects to examine how these factors affect method performance.

To generate realistic expression data, we estimated gene-wise negative binomial (NB) means and dispersions from the human Airway bulk RNA-seq dataset [22] and used these estimates to simulate counts with a predefined set of DE genes. We then modified the simulated data to examine different factors affecting method suitability (rows of Figures 2 and 3). First, we varied sample size from 3 to 24 biological samples per group to examine the statistical power of the Wilcoxon rank-sum test at small sample sizes. Second, we introduced excess zeros or high-expression outliers, each at two severity levels, to evaluate the consequences of deviations from the NB assumption. Third, we replaced raw counts with preprocessed expression values to assess methods with different input requirements. Finally, we introduced a batch variable partially confounded with biological condition, with gene-specific batch effects, to evaluate methods that cannot adjust for covariates. The simulation benchmark comprised 14 scenarios, each generated with 10 random seeds, yielding 140 datasets.

**Figure 2:**
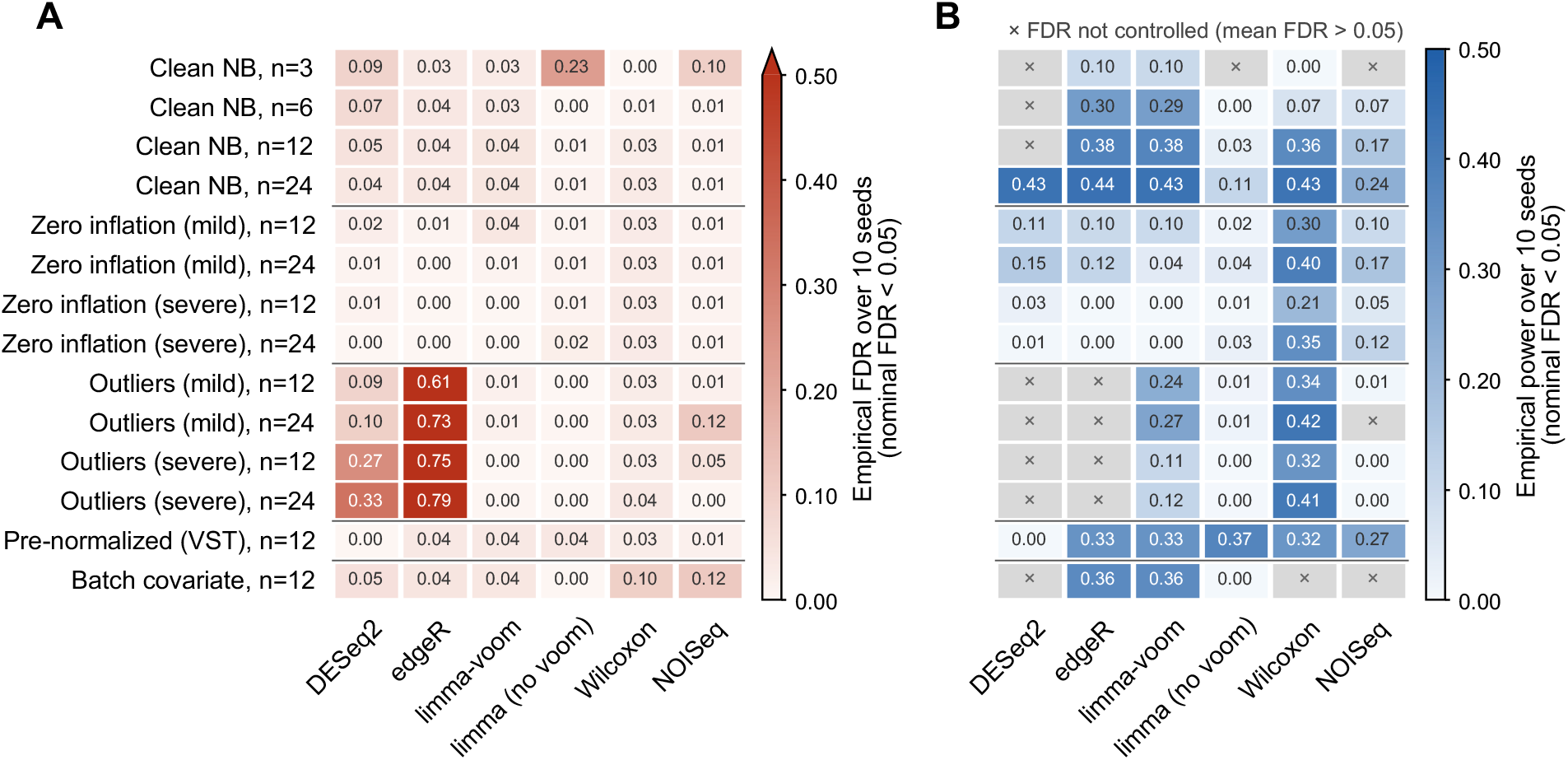
Empirical FDR and power of six DE methods across simulated datasets. (**A**) Empirical FDR at a nominal FDR level of 0.05. (**B**) Empirical power. Values are averaged across 10 simulation replicates generated with different random seeds. Crosses in (B) indicate settings where the empirical FDR exceeded 0.05.

**Figure 3:**
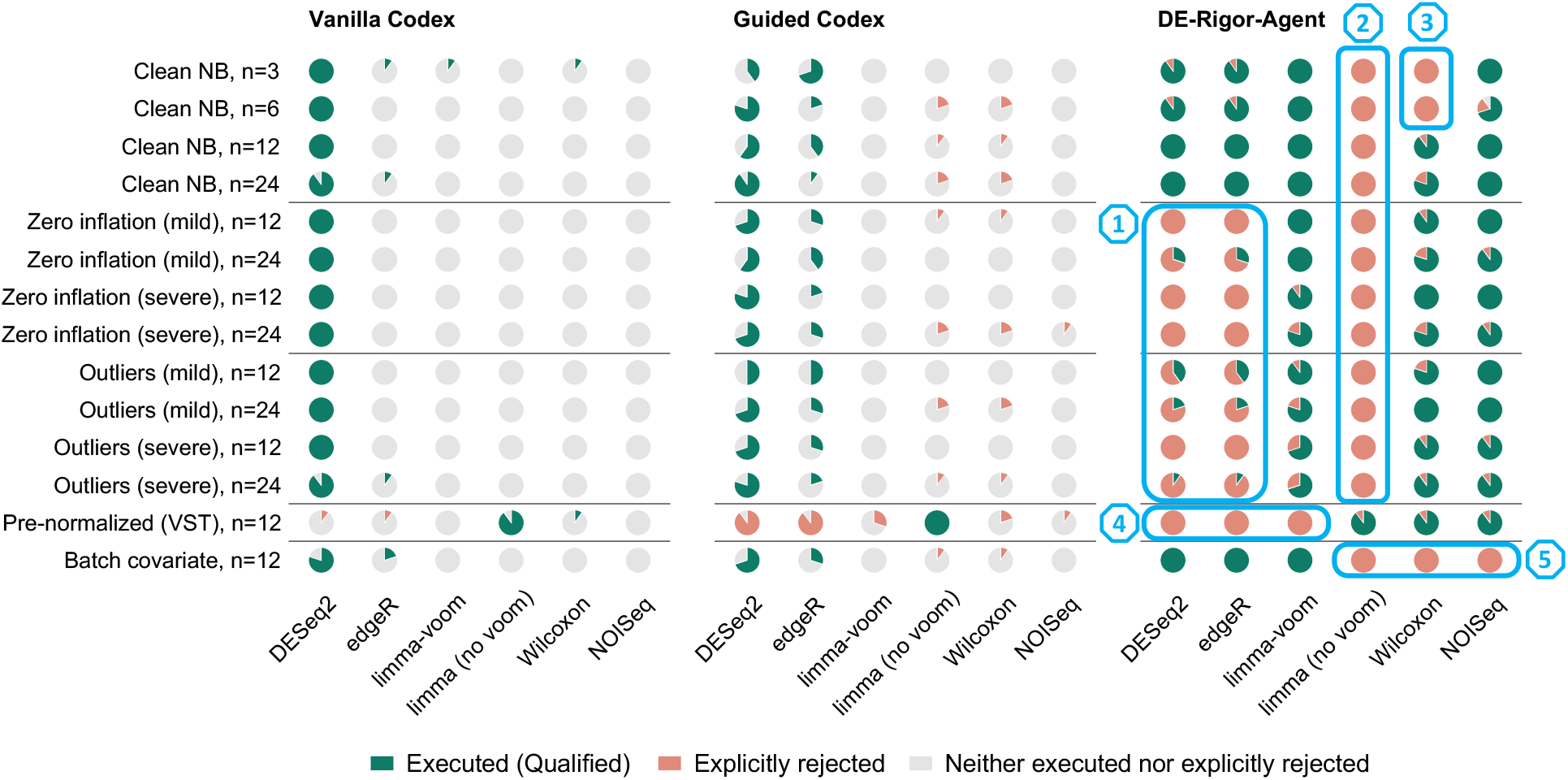
Method-filtering decisions of three agents across simulated datasets. Rows represent simulation scenarios, and columns represent the six candidate DE methods. Each pie chart summarizes 10 simulation replicates generated with different random seeds and shows the proportions of runs in which a method was executed, explicitly rejected with a statistical or data-based rationale, or neither executed nor explicitly rejected. Results are shown for vanilla Codex (left), guided Codex (middle), and DE-Rigor-Agent (right). Guided Codex received an additional instruction to consider method assumptions and data characteristics. Numbered boxes indicate scenario–method combinations targeted for exclusion by the predefined filtering criteria: (1) DESeq2 and edgeR under zero inflation or high-expression outliers; (2) standard limma without voom on raw counts; (3) the Wilcoxon rank-sum test at small sample sizes because of limited statistical power; (4) methods requiring raw counts on pre-normalized input; and (5) methods that cannot adjust for covariates when batch adjustment is required.

We applied all six candidate DE methods to each simulated dataset and evaluated their FDR and statistical power against the known DE status of each gene (Figure 2). On data simulated from NB distributions without additional perturbations, the Wilcoxon rank-sum test had little power with fewer than 12 samples per group. In contrast, edgeR and limma-voom maintained FDR near or below the nominal level while gaining power as sample size increased. DESeq2 also gained power but achieved a mean observed FDR below 0.05 only at 24 samples per group. Deviations from the NB distribution affected method performance differently: excess zeros substantially reduced power without inflating FDR, whereas high-expression outliers led to FDR inflation. The Wilcoxon rank-sum test was comparatively robust to both types of deviations.

Input preprocessing and experimental design also affected method performance. DESeq2 had nearly zero power on pre-normalized inputs, whereas standard limma without voom had low power on raw counts. Under partially confounded batch effects, the Wilcoxon rank-sum test and NOISeq, neither of which supports covariate adjustment in the evaluated implementations, showed inflated FDR. DESeq2 also slightly exceeded the nominal FDR level. Together, these results demonstrate how applying DE methods without considering their statistical assumptions, input requirements, and expected performance can lead to inflated FDR or reduced statistical power, motivating the assumption-aware method filtering implemented in DE-Rigor-Agent.

### 2.3 Evaluation of DE-Rigor-Agent’s Method Filtering

We compared DE-Rigor-Agent with two Codex agent baselines, vanilla Codex and guided Codex, using the 140 simulated datasets. All three agents used the same underlying LLM, GPT-6-Luna, received the same DE analysis request and input data, and considered the same six candidate DE methods. Compared with vanilla Codex, guided Codex received an additional instruction to consider method assumptions and data characteristics. DE-Rigor-Agent followed its structured workflow of input standardization, method filtering, and method execution. None of the agents received information about how the data were simulated. For each candidate method in each run, we classified the outcome as executed, explicitly rejected with a statistical or data-based rationale, or neither executed nor explicitly rejected (Figure 3).

DE-Rigor-Agent documented a decision for every candidate method in all runs, and its exclusions largely matched those expected from the simulation settings (Figure 3, boxes 1–5). In contrast, both Codex agent baselines primarily selected DESeq2 or edgeR, with limited changes in response to the simulated data characteristics. Unlike DE-Rigor-Agent, the baseline agents frequently omitted methods without explicitly explaining why they were not selected.

Vanilla Codex selected DESeq2 for nearly all datasets and rarely explained why other methods were not used. Guided Codex primarily selected DESeq2 or edgeR and occasionally rejected standard limma without voom or the Wilcoxon rank-sum test. However, its method choices changed little across datasets simulated from NB distributions, datasets with zero inflation, and datasets with high-expression outliers, despite the reduced power or inflated FDR observed for some methods in these scenarios (Figure 2). Neither baseline agent systematically excluded NB-based methods when the data deviated from the NB assumption. Both baseline agents selected standard limma without voom for pre-normalized inputs, indicating some sensitivity to input data transformations. In contrast, DE-Rigor-Agent adjusted its filtering decisions according to the data characteristics.

It excluded DESeq2 and edgeR under zero inflation and high-expression outliers when diagnostics indicated deviations from the NB assumption. It excluded the Wilcoxon rank-sum test at small sample sizes because of its limited expected power, while retaining it at larger sample sizes. It also excluded standard limma without voom on raw counts and methods requiring raw counts on pre-normalized inputs. When batch adjustment was required, it excluded methods that could not accommodate covariates. These exclusions were consistent with the predefined filtering criteria and, in many scenarios, avoided methods that showed inflated FDR or reduced power in Figure 2.

These results demonstrate that DE-Rigor-Agent systematically evaluates method suitability using statistical criteria and dataset-specific diagnostics, whereas the Codex agent baselines did not consistently perform these checks, even when explicitly prompted to consider method assumptions. Although filtering does not guarantee optimal FDR control or statistical power among retained methods, it provides a transparent and systematic approach to excluding methods that are unsuitable for the data.

### 2.4 Consistency of Filtering Decisions and Cost

DE-Rigor-Agent’s filtering decisions were largely consistent across simulation replicates. For 77 of the 84 scenario–method combinations, at least 8 of the 10 replicates produced the same decision, and 50 combinations yielded identical decisions across all 10 replicates. Most inconsistencies occurred under mild deviations from the NB assumption, where the diagnostic evidence was less pronounced. For example, DESeq2 and edgeR were retained in 3 of 10 runs with mild zero inflation at 24 samples per group and in 4 of 10 runs with mild outliers at 12 samples per group.

Nevertheless, some retained methods showed inflated empirical FDR relative to the nominal level of 0.05. NOISeq was retained in all runs at 3 samples per group and under mild outliers at 24 samples per group, where its empirical FDR was 0.10 and 0.12, respectively. DESeq2 was retained in 9 of 10 runs at both 3 and 6 samples per group, where its empirical FDR was 0.09 and 0.07, respectively. These results reflect the conservative exclusion strategy of DE-Rigor-Agent: methods are retained unless the predefined criteria identify a violation of their assumptions or input requirements, or a specific performance concern. This approach avoids excluding methods without sufficient justification, but retaining a method does not guarantee its FDR control or statistical power. DE-Rigor-Agent aims to identify methods unsuitable for a given dataset rather than select the single best-performing method.

The method-filtering stage and execution of multiple methods increased the cost of each analysis. We estimated LLM usage costs from the input and output token counts of each run, using the corresponding token prices published by OpenAI. The mean estimated cost per dataset was $0.018 for DE-Rigor-Agent, compared with $0.004 for vanilla Codex and $0.005 for guided Codex (Figure 4). Thus, the more systematic evaluation of method suitability incurred a small additional cost in absolute terms.

**Figure 4:**
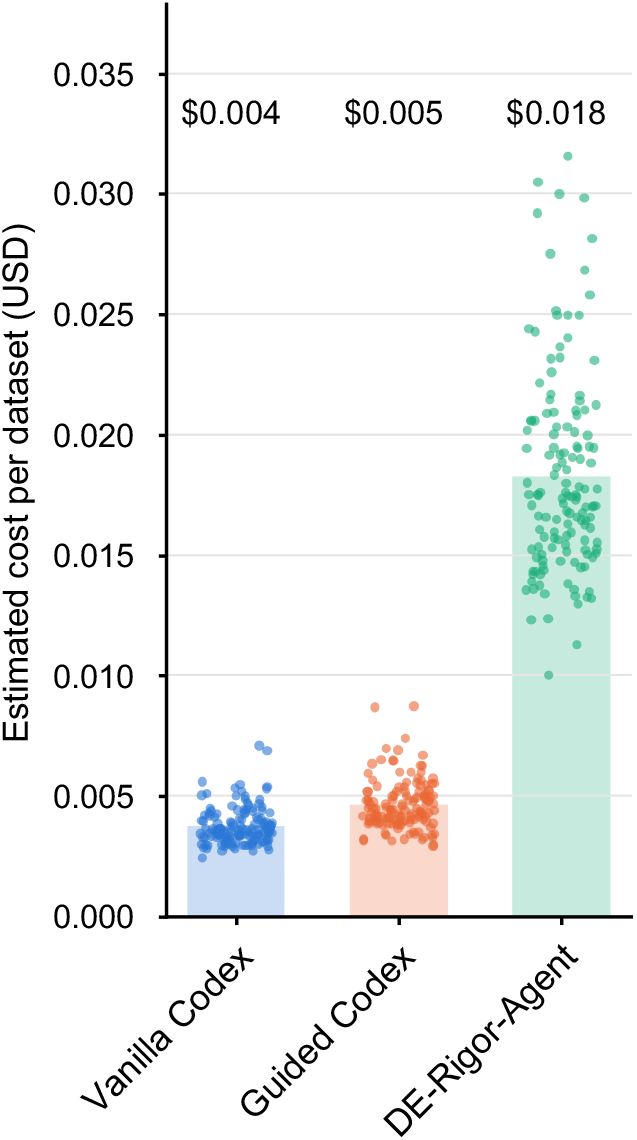
Estimated LLM cost per dataset, calculated from token usage and OpenAI’s published token prices. Bars indicate means across 140 datasets, and points represent individual runs.

## 3 Discussion

We developed DE-Rigor-Agent to improve the statistical rigor and transparency of automated bulk RNA-seq DE analysis through an explicit method-filtering stage. Across 140 simulated datasets, DE-Rigor-Agent systematically evaluated six candidate methods, and its filtering decisions largely matched the expected exclusions based on statistical assumptions, input requirements, and known performance limitations. In contrast, general-purpose Codex agents primarily selected DESeq2 or edgeR without systematically evaluating their suitability for the data, even when explicitly prompted to consider method assumptions. These findings demonstrate the importance of incorporating expert-curated statistical criteria and dataset-specific diagnostics into agent workflows, rather than relying on general-purpose LLM reasoning alone. By evaluating all candidate methods before execution and reporting results from all retained methods, DE-Rigor-Agent also provides a transparent record of method selection and reduces the potential for post hoc selection based on observed DE results.

DE-Rigor-Agent has several limitations. First, method filtering depends on the ability of data diagnostics to detect violations of statistical assumptions. Mild deviations may be difficult to identify, leading to variation in filtering decisions across simulation replicates. Second, satisfying the prede-fined filtering criteria does not guarantee FDR control or high statistical power, as demonstrated by some retained methods in our simulations. DE-Rigor-Agent takes a conservative approach to exclusion: it retains methods unless the available evidence supports their exclusion, rather than attempting to identify the single best-performing method. Finally, our evaluation covered 14 simulated scenarios involving six commonly used DE methods. Further evaluation across additional simulation scenarios and real-world RNA-seq datasets would help establish the framework’s generalizability.

An advantage of DE-Rigor-Agent is that its expert-curated method cards and filtering principles are expressed in natural language, allowing domain experts to review, update, and extend them without changing the underlying agent architecture. The workflow can also accommodate different analysis goals while preserving the predefined statistical criteria for method filtering. More broadly, this design could be extended to other bioinformatics tasks in which selecting appropriate statistical methods requires evaluating their assumptions against dataset characteristics. By incorporating statistical expertise directly into automated workflows, DE-Rigor-Agent provides a practical approach to making computational analyses more rigorous and transparent.

## 4 Methods

### 4.1 DE-Rigor-Agent Implementation and Agent Configurations

DE-Rigor-Agent was implemented in Python using LangChain and LangGraph. Its workflow consists of three sequential stages, implemented as nodes in a fixed graph: input standardization, method filtering, and method execution. At each stage, the LLM follows stage-specific instructions and iteratively calls tools until it produces a structured report. Two tools are available across all stages: a code-execution tool that runs Python or R code in the working directory, and a question tool that requests clarification from the user when necessary. The method-execution stage has an additional tool for launching dedicated subagents.

Each subagent is a separate agent instance using the same LLM as the main agent. It receives a method-specific system prompt containing a complete, executable code template and uses the code-execution tool to adapt and run the template. The subagent then returns a summary of its DE results to the main agent. At the end of each stage, only the stage-specific instructions and final report are retained in the message history; intermediate code and tool outputs are removed from the LLM context. This design limits the information passed between stages while preserving their outputs. The method cards and filtering principles are specified in natural language in the main agent’s system prompt and can be updated without modifying the underlying code. DE-Rigor-Agent supports both a Chainlit web interface and a command-line interface; all experiments in this study used the latter.

To ensure a consistent comparison, DE-Rigor-Agent, vanilla Codex, and guided Codex used the same underlying LLM (GPT-6-Luna) with medium reasoning effort through the OpenAI API, the same Python and R environments, and the same six candidate DE methods. Each run started in a fresh working directory containing only the input dataset table. Scenario labels, simulation parameters, and the true DE status of genes were not provided to any agent.

*Vanilla Codex* (Codex CLI version 0.159.2) received the user prompt “Perform DE analysis on the given data.”, together with the list of six candidate methods and the paths to the Python and R environments.

*Guided Codex* used the same configuration as vanilla Codex, with one additional sentence in the user prompt: “Consider the assumptions of different DE methods and the characteristics of the data to select a statistically rigorous and appropriate method.”

*DE-Rigor-Agent* received the same basic user prompt, “Perform DE analysis on the given data.”, through its command-line interface. Unlike the baseline agents, its six candidate methods and corresponding statistical criteria were predefined in the system prompt.

### 4.2 Simulation Design

We generated baseline gene-expression counts from a negative binomial (NB) generalized linear model:

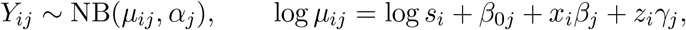

where *i* indexes samples, *j* indexes genes, 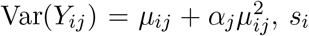 is the library-size factor, *β*_0*j*_ is the baseline log mean, *x*_*i*_ ∈{0, 1} indicates condition, *β*_*j*_ is the log fold change, and *z*_*i*_ ∈ {0, 1} indicates batch with gene-specific effect *γ*_*j*_. Batch effects were included only in the batch scenario; otherwise, *γ*_*j*_ = 0.

Gene parameters were calibrated using the Airway bulk RNA-seq dataset [22]. After removing genes with zero total counts, an intercept-only DESeq2 model yielded 33,469 pairs of base mean and dispersion estimates. We sampled 2,000 genes with replacement, setting *β*_0*j*_ to the log base mean and *α*_*j*_ to the estimated dispersion. Library-size factors were log-normally distributed (log-scale SD = 0.5) and normalized to a geometric mean of 1. Of the 2,000 genes, 400 (20%) were DE: 200 upregulated (fold change 1.6) and 200 downregulated (fold change 0.6). For a given seed and sample size, scenarios shared gene parameters, DE gene assignments, and library-size factors; scenarios with post-simulation modifications also shared the underlying NB counts.

We constructed 14 scenarios: *Clean NB data*: 3, 6, 12, or 24 samples per group, without modifications. *Zero inflation*: at 12 or 24 samples per group, counts were independently set to zero with gene-specific probabilities 0.02 + 0.18(1− *r*_*j*_) (mild) or 0.05 + 0.35(1− *r*_*j*_) (severe), where *r*_*j*_ is the percentile rank of the gene’s base mean. *High-expression outliers*: at 12 or 24 samples per group, observations were independently selected with probability 0.02 (mild) or 0.05 (severe) and replaced by ⌈*m* max(*Y*_*ij*_, *q*_*j*_) ⌉, where *q*_*j*_ is the gene’s upper-quartile count (at least 1) and *m* ∼*U* (10, 25) (mild) or *U* (20, 50) (severe). Zero inflation and outliers were introduced independently of condition and DE status. *Pre-normalized input* : at 12 samples per group, counts were transformed using 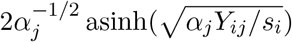, retaining non-integer values. *Confounded batch effects*: at 12 samples per group, batch A/B composition was 9/3 in controls and 3/9 in treatment. An additional 400 non-DE genes received batch-B effects (200 with fold change 1.6 and 200 with fold change 0.6), without condition-by-batch interactions.

Each scenario was simulated with 10 random seeds, yielding 140 datasets. Agents received a single table containing sample identifiers, conditions, batch annotations when applicable, and gene-expression values.

### 4.3 DE Method Implementation and Performance Evaluation

We applied all six candidate DE methods to every simulated dataset, regardless of method suitability, using R 4.4.3 (DESeq2 1.46.0, edgeR 4.4.2, limma 3.62.2, NOISeq 2.50.0). In the batch scenario, DESeq2, edgeR, limma-voom, and standard limma used the design `∼batch+condition` and tested the condition effect; the Wilcoxon rank-sum test and NOISeq were run without batch adjustment.

Method implementations were as follows. *DESeq2* : genes with at least 10 counts in at least *n* samples (*n* = samples per group), followed by the default `DESeq` workflow and Wald test; pre-normalized values were rounded, and gene-wise dispersions were used when dispersion-trend fitting failed. *edgeR*: `filterByExpr`, TMM normalization, and quasi-likelihood F-test (`glmQLFit, robust = TRUE`). *limma-voom*: filterByExpr, TMM normalization, voom, and eBayes (`robust = TRUE`). *Standard limma*: lmFit and eBayes (robust = TRUE) applied directly to supplied values without normalization or transformation. *Wilcoxon rank-sum test* : `filterByExpr`, TMM-normalized counts per million (CPM), and two-sided normal-approximation tests. *NOISeq* : `noiseqbio`, TMM normalization, 50 permutations, *k* = 0.5, and internal CPM filtering.

Genes were called DE at Benjamini–Hochberg (BH)-adjusted *p <* 0.05, except for NOISeq, which used a differential expression probability threshold of 0.95 [8]. For each dataset, the false discovery proportion (FDP) was the fraction of called genes that were truly non-DE (zero if none were called), and power was the fraction of the 400 true DE genes detected, counting filtered-out genes as undetected. Empirical FDR and power were calculated by averaging FDP and power across 10 simulation replicates. Crosses in Figure 2 indicate settings where empirical FDR exceeded 0.05.

### 4.4 Evaluation of Method-Filtering Decisions

For each agent run, we classified the outcome for each of the six candidate DE methods into three categories. A method was classified as *Executed* if its DE analysis completed successfully and the results were reported. Using a method only for preprocessing or diagnostics, such as applying DESeq2’s variance-stabilizing transformation, did not count as execution. A method was classified as *Explicitly rejected* if it was not executed and the agent provided a statistical or data-based reason for excluding it. Simply preferring another method or abandoning execution after a software error did not count as explicit rejection. All remaining cases were classified as *Neither executed nor explicitly rejected*, including methods omitted without explanation or never mentioned by the agent. This third category occurred only for the two Codex agent baselines, as DE-Rigor-Agent explicitly retained or rejected every candidate method.

We determined these outcomes from each run’s generated code, execution records, and final response, together with the method-filtering report for DE-Rigor-Agent. An independent LLM auditor (Claude Sonnet 5.5) reviewed each run and classified the outcome for every candidate method, providing supporting quotations from the analysis records. These classifications were checked against the recorded evidence and cross-validated using rule-based detection of executed methods from the generated code and execution records. For DE-Rigor-Agent, explicit rejections were additionally verified against its method-filtering report.

We estimated the LLM cost of each run from its token usage and the published GPT-6-Luna token prices. The rates per million tokens were $0.10 for uncached input, $0.01 for cached input reads, $0.125 for cache writes, and $0.50 for output. Total cost summed these token costs across all LLM calls in each run. Because Codex does not report cache-write tokens separately, we charged at the uncached-input rate, which may underestimate its cost by up to 18% per run.

## Code Availability

The source code, all agent prompts, and the expert-curated knowledge of DE-Rigor-Agent are available at https://github.com/zqrong/de-rigor-agent.

## Notes

### Competing Interest Statement

The authors have declared no competing interest.

